# RNA order regulates its interactions with zwitterionic lipid bilayers

**DOI:** 10.1101/2024.08.28.610126

**Authors:** Akhil Pratap Singh, Janak Prabhu, Stefano Vanni

## Abstract

RNA-lipid interactions directly influence RNA activity, which plays a crucial role in developing new applications in medicine and biotechnology. However, while specific preferential behaviors between RNA and lipid bilayers have been identified experimentally, their molecular origin remains unexplored. Here we use molecular dynamics simulations to investigate the interaction between RNA and membranes composed of zwitterionic lipids at the atomistic level. Our data reproduce and rationalize previous experimental observations, including that short-chain RNAs rich in guanine have higher affinity for gel-phase membranes compared to RNA sequences rich in other nucleotides and that RNA prefers gel-phase membranes to fluid bilayers. Our simulations reveal that RNA order is a key molecular determinant of RNA-zwitterionic phospholipid interactions. Our data provides a wealth of information at the atomic level that will help accelerating research on RNA-lipid assemblies for task-specific applications such as designing lipid-based nanocarriers for RNA delivery.

## Introduction

RNA oligomers play a crucial role in controlling several cellular functions, including cell death and proliferation^1^, and they are associated with the onset of several types of cancer and other genetic disorders in cells.^2–5^ As a consequence, RNA-based therapeutics have experienced significant progress in the last few years, leading to a breakthrough in gene therapy approaches for the treatment of a wide range of diseases.

RNA requires robust delivery systems to prevent the loss of RNA activity^6^, and phospholipid-RNA conjugates have shown great promise as delivery carriers^4,7–10^. Yet, how lipids interact with various RNA-based therapeutics, such as messenger RNA (mRNA) or small interfering RNA (siRNA) at the molecular level remains mostly unclear.

Experimental studies have demonstrated that nucleotide sequence and structure greatly influence molecular interactions in phospholipid-RNA conjugates.^11–19^ For example, in lipid nanoparticles, siRNA resides on the surface of conjugates and mRNA at its centre.^11–13^ Specifically, such interactions are strongly influenced by electrostatics, since RNA has a negative charge on the phosphates (PO_4_ ^−^) and numerous cellular lipids have a variety of differently charged head groups,^18^ including lipids with negatively to zwitterionic charged moieties.^19^

In addition to electrostatic interactions, several reports have shown that RNA/DNA adsorption at the bilayer interface can also be controlled by acyl chain order ^20–28^, with randomized mixtures of RNA showing a preferential binding affinity for membranes in the solid crystalline (gel) phase.^20,26^ Moreover, recent studies have revealed that specific RNA sequences (guanine rich) display higher binding affinity for gel membranes in the presence of low salt concentrations.^22^ Additionally, it has been reported that RNA–lipid binding regulates RNA activity^29,30^ and that lipids can provide RNA-safeguarded and selective micro compartments^31,32^ through membranous structures due to their amphiphilic nature; specifically, RNA could reside on membrane surfaces as a consequence of direct RNA-lipid interactions providing a specialized physicochemical microenvironment to maintain RNA activity.^30^

However, none of the studies described above, presumably due to the inherent limitations involved in the techniques used, provided a detailed explanation for the molecular origin of these specific RNA-zwitterionic lipid interactions. To investigate this aspect, here we opted to use all-atom (AA) molecular dynamics (MD) simulations, as this computational technique can provide an atomic-level description of complex chemical systems under highly controlled conditions, complementary to experimental methods. By focusing on the detailed understanding of the RNA– lipid binding as a function of the primary sequence, structure, and length of RNA as well as on the nature of the membrane (ordered *vs* liquid phase), we found that RNA order is the key property modulating binding of RNA to zwitterionic phospholipid bilayers. Our study provides useful information at the molecular-to-atomic level, which could be helpful to design robust lipid nanoparticles as efficient drug delivery systems.

## Results and Discussion

### Guanine rich oligomers exhibit greater adsorption at DPPC-gel phase bilayers

To investigate the molecular origin of the interactions between RNA and lipid bilayers, including the puzzling specificity for guanine observed experimentally^22^, we initially performed AA-MD simulations of RNA oligomers, with variations in their sequence and structure, on DPPC-gel phase systems. We simulated four different short-chain oligomers, with each system containing a homo-oligomer composed of a four-nucleotide sequence (4-nt): adenine (A4), cytosine (C4), guanine (G4) and uracil (U4) (Fig. 1A). These simulated systems and their compositions are summarized in Table S1, and method details can be found in Supplementary Information.

**Figure 1.**
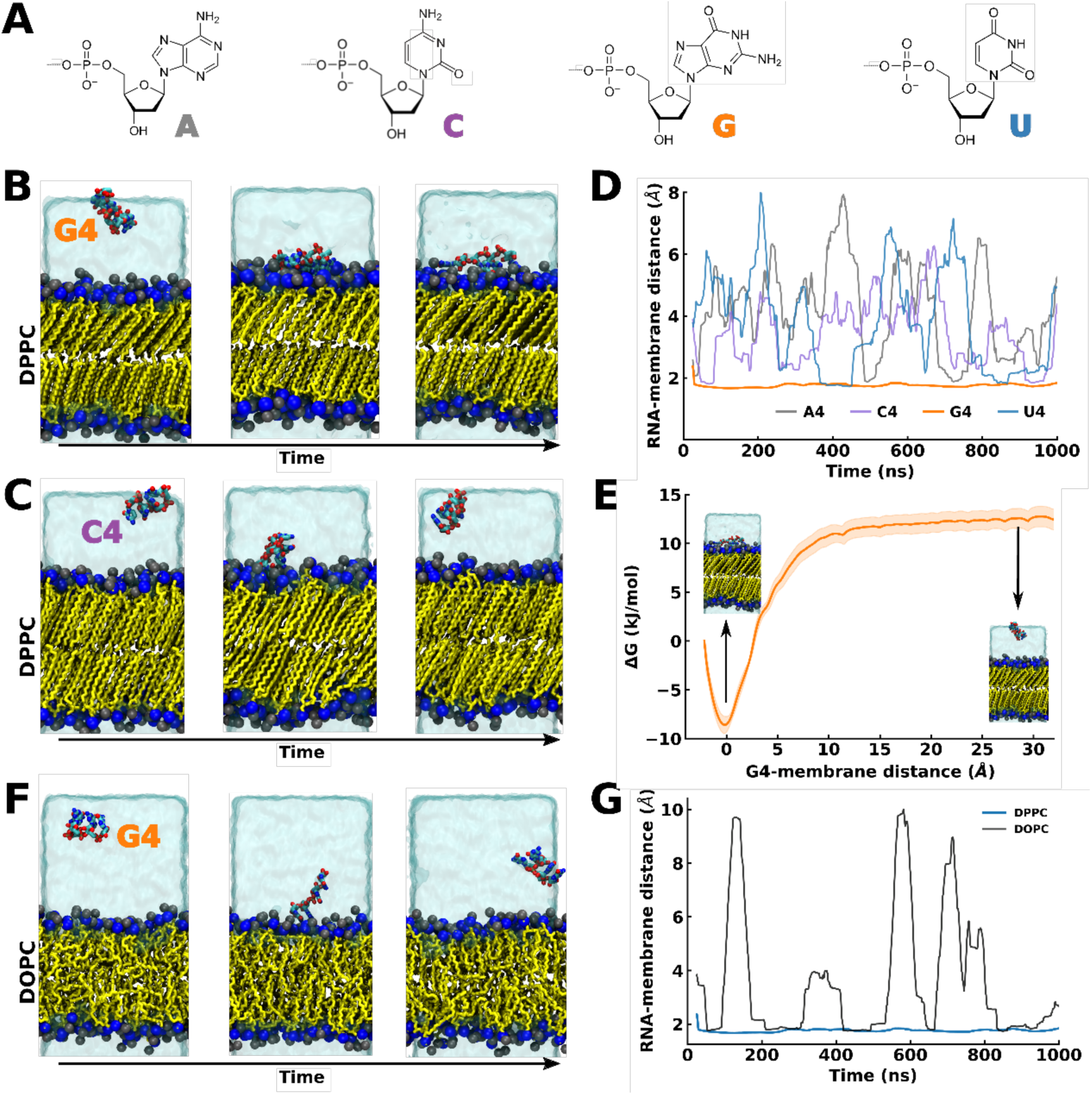
G4, but not A4, C4 and U4, stably binds to DPPC-gel lipid bilayers. *(A)* Molecular structures of RNA nucleobases: adenine, cytosine, guanine and uracil. *(B, C)* Representative snapshots of the time evolution of MD simulation of *(B)* G4 oligomer/DPPC-gel bilayer, and *(C)* C4 oligomer/DPPC-gel bilayer systems. Color scheme: Blue and black for nitrogen and phosphate atom of lipid head groups, yellow for lipid tails, translucent ice blue for water. Oligomers (G4 or C4) are in multicolored licorice representation. Counter ions are removed for clarity. *(D)* Running average of the minimum distance between oligomers and the DPPC-gel bilayer over simulation time, *(E)* Potential of mean force (PMF) for G4 oligomer binding to the bilayer surface along the bilayer normal (z-axis), with snapshots of bound and unbound state from the umbrella sampling simulations. *(F)* Representative snapshots of the time evolution of MD simulation of G4/DOPC-liquid bilayer system. The color scheme is identical to that of Fig. 1B. *(G)* Running average of the minimum distance between oligomers and lipid bilayer in G4/DPPC-gel bilayer and G4/DOPC-liquid systems.

Analysis of the MD trajectories reveals that guanine-rich oligomers (G4) easily adsorb on bilayer interfaces (Fig. 1B, D; S1, S2). G4 adsorption is maintained during the time scale of our simulations (Fig 1D). In contrast, the interaction of C4, A4 and U4 oligomers with the interface is short-lived or absent (Fig. 1C, D; S1, S2), as these oligomers do not create stabilizing interactions at the interface, quickly returning to the aqueous phase (Fig. 1C, D).

To characterize the preference of the different oligomers for the DPPC-gel bilayer, we first computed the interaction energy (*E_Coul._* + *E_LJ_*) between the different oligomers and the DPPC-gel bilayer over time (Fig. S1C). This analysis indicates that the interaction between G4 and the bilayer has a strong enthalpic component, achieved via the establishment of electrostatic and Lennard Jones interactions between the oligomer and the lipid head groups. Next, to explicitly quantify the binding free energy of G4 to the DPPC-gel bilayer (Fig. 1E), we performed umbrella sampling calculations using the distance between the COM of G4 and the bilayer as the reaction coordinate. This analysis also identifies a clear minimum at the bilayer interface, with a binding free energy for G4 of approximately 22 kJ/mol (Fig. 1E).

Overall, all these analyses indicate that G4 mainly localizes at the water-bilayer interface, while all other oligomers (C4, A4, U4) mostly reside in water (Figs 1 and S1). Of note, control simulations with physiological ion concentrations and in the presence of divalent ions (150 mM NaCl + 50mM MgCl_2_), indicate that these results are qualitatively similar in the presence of physiological ionic conditions, with G4 stably binding DPPC-gel bilayers more readily than A4. However, a moderate binding of A4 to the DPPC-gel bilayer at this ionic strength, which is significantly higher than that used experimentally^22^, could be observed (Fig. S2).

Finally, we investigated the interaction of G4 with fluid DOPC bilayers, as short nucleotide oligomers were shown experimentally to strongly prefer gel-phase *vs* fluid-phase bilayers^22^ (Fig. 1F, G). In agreement with experimental observations, we indeed observed significantly lower membrane binding to DOPC bilayers for G4, as indicated by the time trace of the minimum distances between G4 and the bilayer along the MD trajectory and by G4-bilayer distances probability distribution with time (Fig. 1G and Fig. S3). Overall, our data indicate that nucleotide oligomers do not interact with fluid DOPC bilayers, and that G4 establishes preferential interactions with DPPC-gel bilayers compared to all other nucleobases, in agreement with experimental measurements.^22^

### Base-pairing promotes binding of RNA to gel-phase zwitterionic lipid bilayers

We next investigated larger-scale RNA structural effects on RNA-bilayer binding, and specifically the decreased propensity of single-stranded RNA (ssRNA) *vs* double-stranded RNA (dsRNA) to bind to DPPC-gel bilayers observed experimentally.^22^ To this end, we simulated three 24-nt ssRNAs—one cytosine/guanine (GC)-rich, one uracil and one adenine homo-oligomers—and one dsRNA (Fig. 2A) with a DPPC-gel bilayer. We took three different ssRNA as they exhibit sequence-dependent ordered and disordered conformations.^33, 34^ Specifically, both the adenine homo-oligomer and the cytosine/guanine (GC)-rich are expected to form ordered structures due to favorable base-pairing or stacking interactions, while the uracil homo-oligomer is expected to remain unfolded.^33^ For dsRNA, we simulated multiple dsRNA structures obtained from the PDB in an aqueous environment to confirm their duplex stability over time. After proper screening, dsRNA from PDB ID 4JRT was chosen as a model for these simulations. System details are given in Table S1, while Table S2 lists the RNA sequences used.

**Figure 2.**
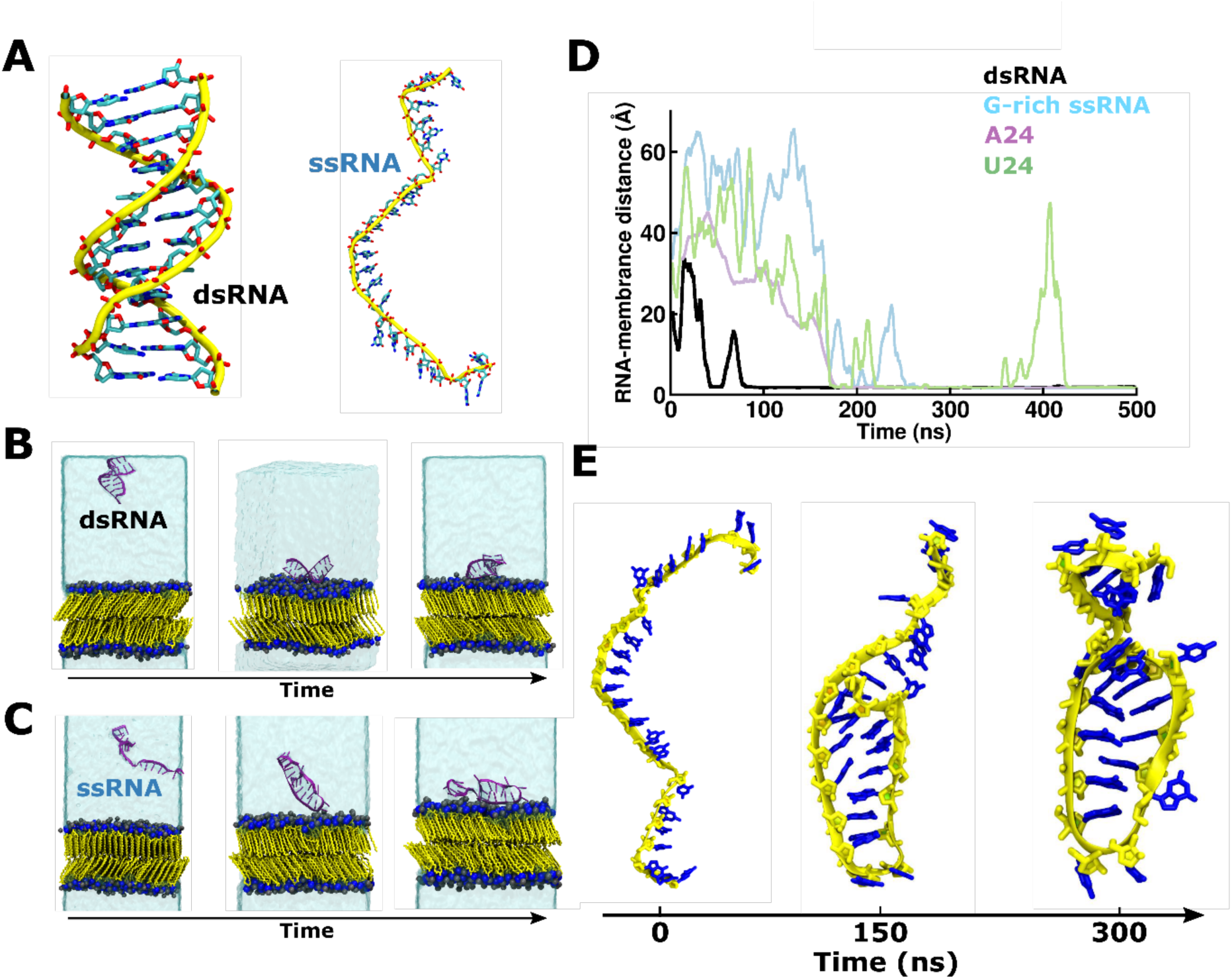
dsRNA and ssRNA adsorption on DPPC-gel membranes. *(A)* Structure of simulated (i) dsRNA and (ii) ssRNA. *(B-C)* Time evolution of MD simulations of *(B)* dsRNA/DPPC-gel, and *(C)* G-rich ssRNA/DPPC-gel systems. Color scheme: Blue and black for nitrogen and phosphate atom of lipid head groups, yellow for lipid tails, translucent ice blue for water. RNA structures are depicted in new-cartoon (magenta) representation. *(D)* Time evolution of the minimum distance between dsRNA or ssRNA and DPPC-gel bilayer in MD simulations. *(E)* Structural arrangement of G-rich ssRNA in ssRNA/DPPC-gel phase bilayer system over time, at (i) t = 0ns, (ii) t = 150 ns and (iii) t = 300 ns. Color scheme: yellow for ssRNA backbone and blue for nucleic bases.

Analysis of the MD trajectories of both dsRNA/DPPC and ssRNA/DPPC systems indicates that all systems adsorb on the membrane interface (Fig. 2B, C, D). This observation is consistent with experimental observations,^22^ as the RNA/lipid ratio in our *in silico* systems (12 lipids per nucleotide) is above the threshold at which membrane binding is experimentally observed for both dsRNA and ssRNA.^22^ However, analysis of the time-traces indicates that dsRNA always binds to the DPPC-gel bilayer within a few tens of nanoseconds of simulation, staying there for the rest of the simulation (Fig. 2B, D; S4), while binding of ssRNA is delayed compared to dsRNA (Fig. 2C, D).

Further, analysis of the trajectories reveals that in all our replicas, G-rich and adenine-homo-oligomers (Fig. S5A, B and Fig. S6A, B) switch to a stable, ordered structure (comparable to that of a hairpin) in a sequence-dependent manner,^33, 34^ somewhat analogous to dsDNA, before adsorption on the bilayer (Fig. 2C, E, and Fig. S7A, B). On the contrary, U24-ssRNA, in agreement with experimental observations^33^ displays a persistent disordered structure and long-delayed adsorption in all replicas (Fig. S8A, B, C).

To investigate the molecular origin of the different dsRNA and ssRNA-U24 adsorption mechanisms, we focused on the two major interactions driving RNA binding to lipid bilayers: (i) charge-charge interactions between the negatively charged phosphate groups on the oligomer backbone and the positively charged moieties of lipid head groups, and (ii) hydrogen bonds between the nucleotide side chains and the phospholipid head groups. We found that charge-charge interactions are the dominant energy term for the initial RNA-membrane adsorption (Fig. S9A) and that while direct H-bonds between dsRNA and the lipid bilayer are almost absent throughout the trajectory, H-bond interactions between U24 and the DPPC-gel bilayer become an important contribution in the later stages of U24 adsorption (Fig. S9C), resulting in an increase in enthalpic interaction with the bilayer (Fig. S9B). Overall, our simulations suggest that the mechanism of RNA adsorption correlates with its secondary structure, suggesting that predefined ordered states might preferentially accelerate its binding to DPPC-gel lipid bilayers. As a preliminary confirmation of this hypothesis, we observed that ssRNA binds to bilayers much quicker in the presence of high salt concentrations and divalent cations, as these conditions promote a stabler structure for ssRNA oligomers (Fig S10).^33,35–41^

### RNA order facilitates the interaction between RNA and gel-phase lipid bilayers

The observation that ssRNA sequences such as G-rich and A24 oligomers fold into ordered structures before binding to DPPC-gel bilayers suggests that ssRNA disorder might antagonize its binding to lipid bilayers. To assess the generality of this observation, we next tested whether this holds true also for the simple case of the 4 small RNA oligomers (A4, C4, G4 and U4) we previously investigated, and for which membrane binding was observed exclusively for the oligomer G4 (Fig. 1).

To this end, we focused on the order/disorder behavior of the 4 small RNA oligomers (A4, C4, G4 and U4) in solution. To do so, we computed their conformational entropy by assessing their microstate populations using cluster analysis.^42^ This analysis indicates that while U4 is highly disordered, all other short oligomers (A4, C4 and G4) adopt one dominant population in solution and have low conformational entropies (Fig. 3A, B). However, structural analysis of these conformations indicate that the G4 oligomer adopts a much more extended structure in solution, as indicated by its radius of gyration (Fig. 3C) and its shape anisotropy (Fig. 3D).

**Figure 3.**
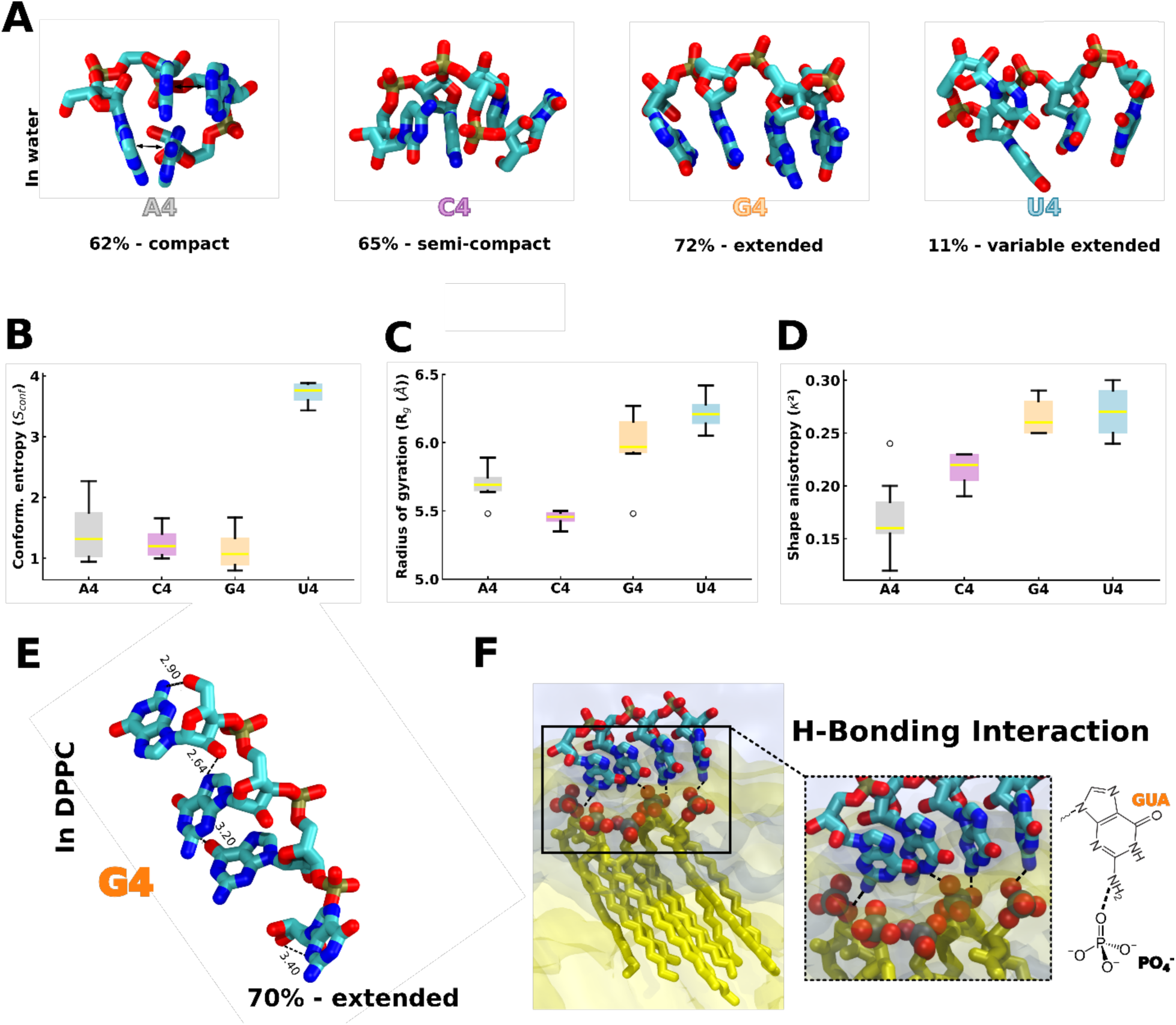
Conformational entropy of RNA oligomers. *(A)* Most populated conformations of the different oligomers from cluster analysis in bulk water phase from the last 600 ns of MD simulations (1000 structures each). *(B)* Oligomers conformational entropy determined from cluster analysis. Color scheme: gray for A4, purple for C4, orange for G4 and blue for U4 oligomers. *(C)* Radius of gyration calculated during the MD simulation for different oligomers in bulk water phase. *(D)* Relative shape anisotropy 𝜅^2^ for all oligomers in bulk water phase. (*E)* Spatial arrangements of the G4 oligomers in G4 oligomer/DPPC-gel system from MD simulations. Color scheme: red for oxygen, cyan for carbon and blue for nitrogen. For clarity, the hydrogens of oligomers are omitted. The dotted lines represent the H-bonds. *(F)* G4 oligomer arrangement at DPPC-gel bilayer interface from MD simulations. The representative MD snapshot of G4/DPPC-gel bilayer showing the probable H-bonding interaction between guanine –NH_2_ group and lipid phosphate oxygen atoms, at interface. Color scheme: red for oxygen, blue for nitrogen, cyan for carbon, brown for phosphate and yellow for lipid tails. Licorice representation (except lipid’s tails) is used for G4 oligomers. The nitrogen, oxygen, and phosphate–atoms of lipid are represented by spheres. The translucent yellow and ice blue represent the DPPC bilayer and water phases. For clarity, only few lipids of DPPC bilayers are shown here and ions are omitted.

Notably, cluster analysis of the G4 oligomer when bound to the DPPC-gel bilayer (Fig. 1) indicates that the same structure that is populated in solution is also the most abundant conformation in the G4-DPPC-gel system (Fig. 3E, F, A). This unique behavior of G4 is likely a consequence of its ability to form multiple H-bonds compared to other nucleotides (Fig. S11). This allows G4 to establish multiple hydrogen bonds with the DPPC-gel bilayer (Fig. 3F, S11) while also retaining multiple intramolecular H-bonds that keep it in an extended conformation (Fig. 3F).

Interestingly, since the area per lipid of the DPPC-gel bilayer is much smaller compared to that of the DOPC bilayer (50 Å^2^ *vs* 68 Å^2^), thus enhancing local surface charge density by 25% in the case of the DPPC-gel bilayer, it appears that the ability of ssRNA to adopt a structure that matches the charge distribution of the bilayer is a key molecular requirement to promote its binding to the membrane.

## Conclusions

Here we used atomistic MD simulations to investigate the adsorption behavior of different RNA structures on zwitterionic lipid membranes and to determine the molecular details of RNA-zwitterionic lipid interactions. While the role of negatively charged lipids and of RNA-lipid electrostatic interactions has been widely investigated in the past, our work emphasizes the binding mechanism of RNA toward gel and fluid-phase lipid bilayers that are devoid of negatively charged lipids. The rationale behind this choice is twofold: not only these interactions have been shown to be important experimentally^22^, but they also could be leveraged to engineer RNA-lipid nanoparticles with targeted physicochemical properties and RNA activity.

In agreement with experimental observations^22^, we observed that guanine-rich short oligomers have higher affinity for DPPC-gel phase membranes compared to other nucelobases. We found that guanine-rich oligomers prefer extended conformation in solution with low conformational entropy, a combination that is achieved thanks to the large number of H-bond acceptor and donor atoms in its molecular skeleton compared to other nucleobases. This facilitates lipid-RNA interaction through more exposed interaction sites and via the matching of the periodicity of polar heads in gel bilayers.

We also observed that dsRNA adsorbs rapidly on DPPC-gel bilayers, likely as a consequence of the perfect matching between the exquisite periodicity of dsRNA that is achieved through base pairing, and that of gel lipid bilayers. Unexpectedly, we also observed that the ability of ssRNA to fold into ordered structures promotes its adsorption on the lipid bilayer. Overall, our results are consistent with previous theoretical considerations suggesting that RNA behaviour is modulated by the combination of electrostatic interactions, hydrogen bonding as well as interactions with ions. ^35–41, 43–45^ However, our simulations also suggest that these interactions not only play a direct role in determining RNA adsorption to lipid bilayers, but they also indirectly modulate this process by strongly affecting RNA conformational plasticity and order/disorder behaviour.

The main limitations of our MD simulations originate from the potential inaccuracy of the employed force field, and from the imperfect matching of the experimental and simulations conditions, e.g. for what pertains RNA/lipid ratio, salt type and concentration as well as insufficient sampling. On the other hand, our simulations provide interesting insights beyond the specifics of RNA-lipid adsorption. First, the excellent agreement of our data with experimental observations on RNA-lipid bilayers interactions validates *a posteriori* the quality of our force fields and simulation parameters, paving the way for future studies of RNA-lipid interactions. Second, our observations raise interesting and practical ideas on how to optimize RNA-lipid interactions for delivery systems− (i) ssRNA binding could be accelerated by increasing the number of nucleobases in the sequence that can do canonical base pairing. Overall, base pairing plays an essential role in adsorption, as we observed in the dsRNA and structured ssRNA in this study. (ii) RNA membrane adsorption, and especially of disordered ssRNA, can be improved by increasing nucleotides that have H-bond donor and acceptor capability, such as guanine or, alternatively, adenine. (iii) Membrane adsorption can be controlled by modifying the order/disorder of the RNA sequence, for example by chemical modifications that alter RNA rigidity. Overall, our study provides valuable information which can be used to expand the current applications of lipid-oligonucleotide conjugates.

## Acknowledgements

The authors gratefully acknowledge Ms. Jennifer Sapia for helping with figure preparation, and Mr. Ashutosh Kumar and Dr. Pablo Campomanes for insightful discussions. This work was supported by the Swiss National Science Foundation (grants CRSII5_189996 to SV). This work was supported by the Swiss National Science Foundation through the National Center of Competence in Research Bio-Inspired Materials. This work was supported by grants from the Swiss National Supercomputing Centre under projects ID s1189 and s1221.

## Supporting Information for

**Figure S1.**
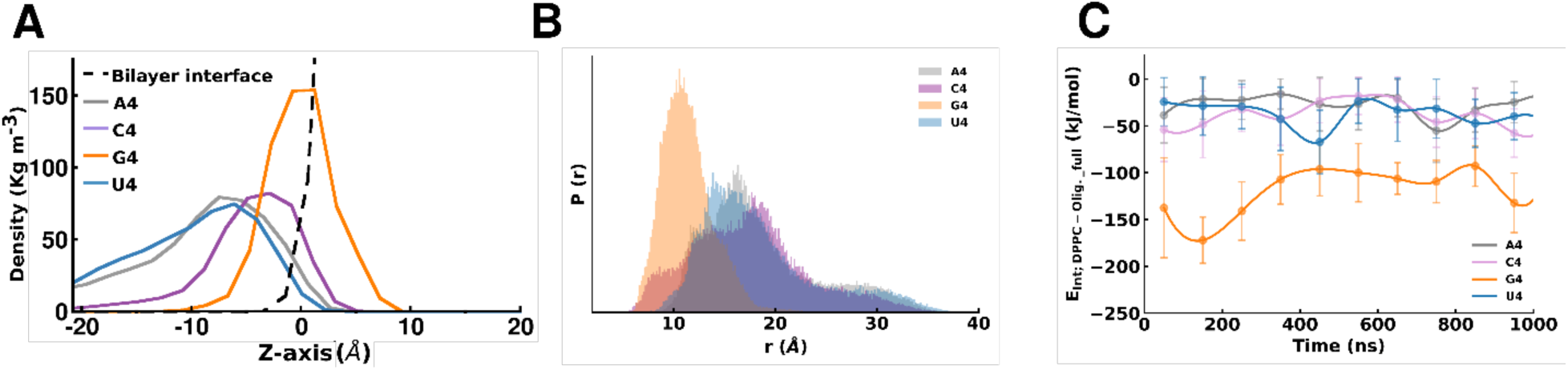
***(A)*** Comparison of different oligomers density profiles along the normal of the DPPC bilayer plane. *(**B**)* Probability distribution of distances between the center of mass (COMs) of oligomers and bilayer head group observed in the complete trajectories (1μs). (***C***) Interaction energy (*E_Coul._* + *E_LJ_*) per nucleotide between the various oligomers and the DPPC-gel bilayer over time. Error bars on the energy curves were estimated using block averaging.

**Figure S2.**
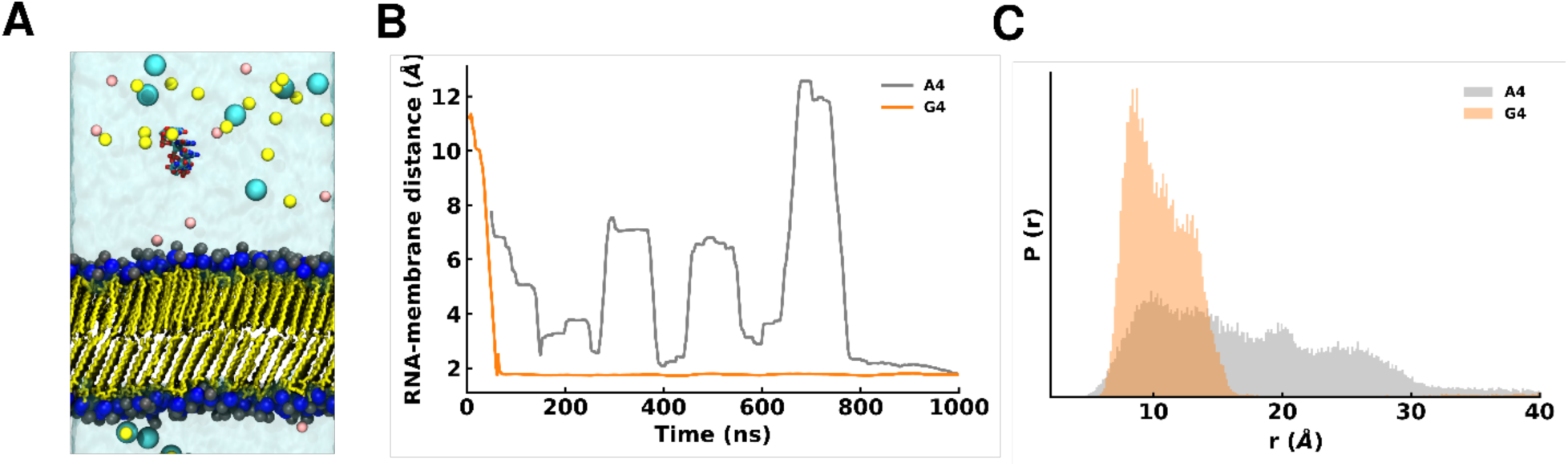
*(A)* Representative snapshot from MD simulation of A4 oligomer/DPPC-gel bilayer system in presence of *150mM NaCl* and *50 mM MgCl_2_* ions. Color scheme: Blue and black for nitrogen and phosphate atom of lipid head groups, yellow for lipid tails, translucent ice blue for water, light yellow spheres for sodium ions, cyan for chloride ions, pink for magnesium ions. The A4 oligomer is in multicolored licorice representation. *(B)* Running average of the minimum distance between oligomers and lipid bilayer in A4/DPPC-gel bilayer and G4/DPPC-gel bilayer systems, and *(C)* probability distribution of distances between the center of mass (COMs) of oligomers and bilayer head group observed in complete trajectory.

**Figure S3.**
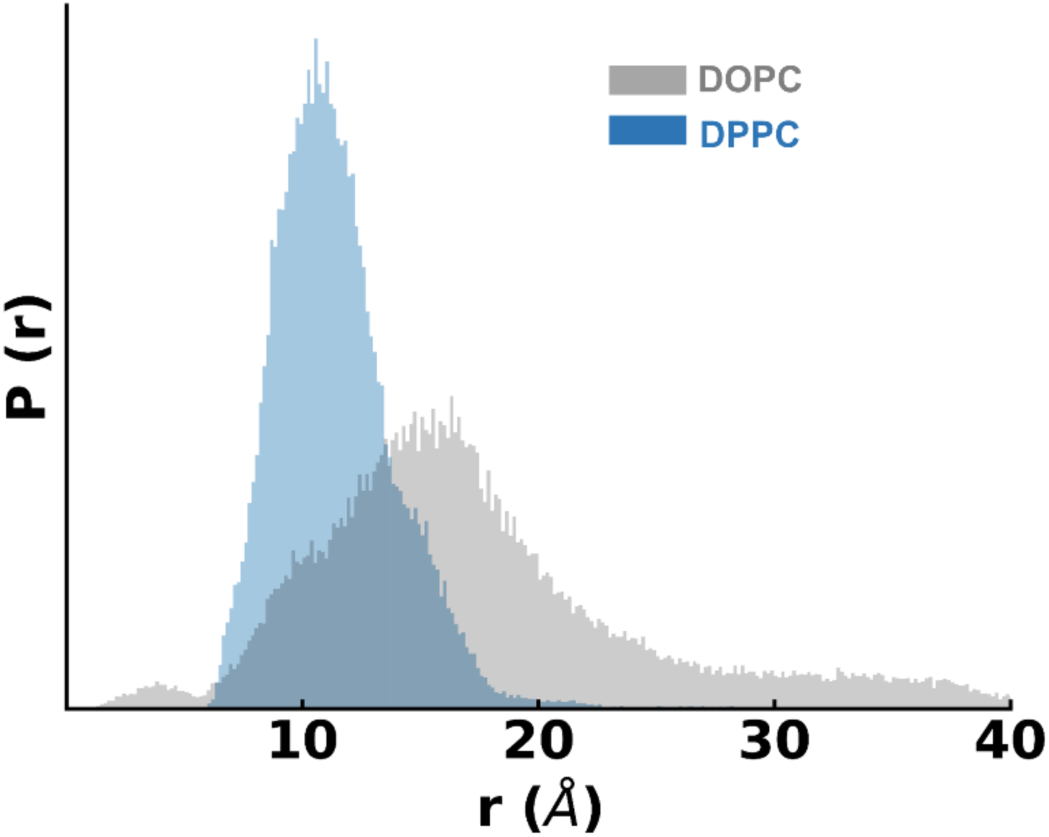
Probability distribution of distances between the center of mass (COMs) of the G4 oligomer and DPPC-gel (blue) and DOPC (gray) bilayer head group observed in the MD simulations.

**Figure S4.**
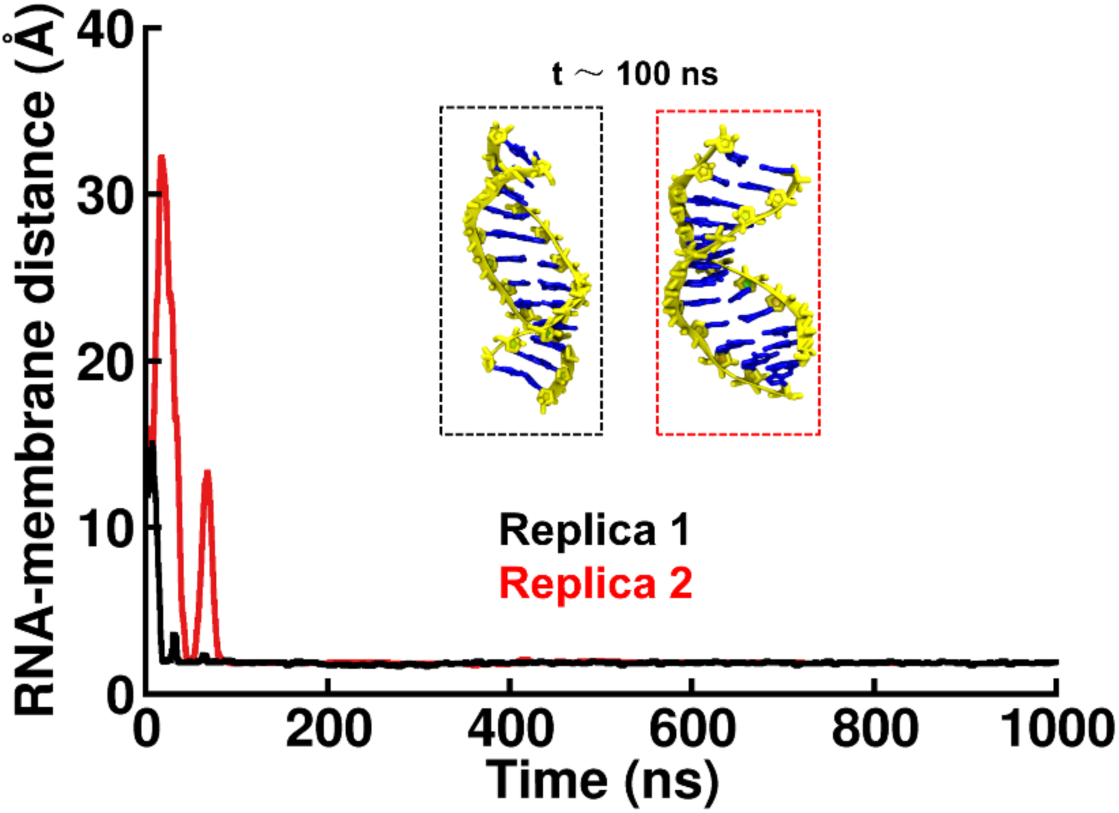
Minimum distance between dsRNA and DPPC-gel bilayer. Inset: structural arrangement of dsRNA in both dsRNA/ DPPC-gel bilayer replicas at t = 100 ns. Both replicas show quick absorption of dsRNA at DPPC-gel bilayer.

**Figure S5.**
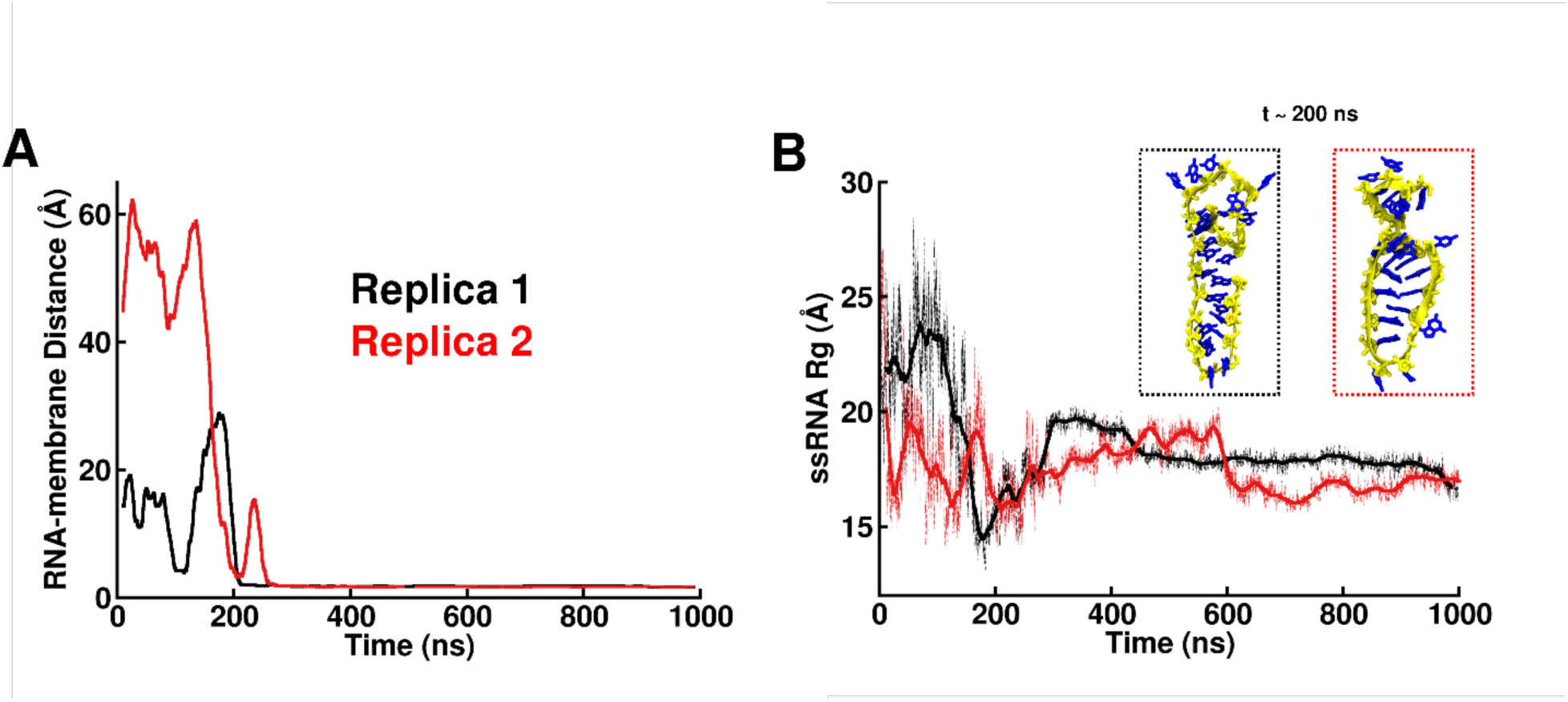
(A) Minimum distance between G-rich ssRNA and DPPC-gel bilayer. (B) Radius of gyration G-rich ssRNA over the simulation time. Inset: structural arrangement of G-rich ssRNA in both ssRNA/DPPC-gel bilayer replicas at t = 300 ns. Fig S5A and Fig S5B show that ssRNA took approximately 200 ns to fold, which concomitantly led to adsorption to the DPPC-gel bilayer.

**Figure S6.**
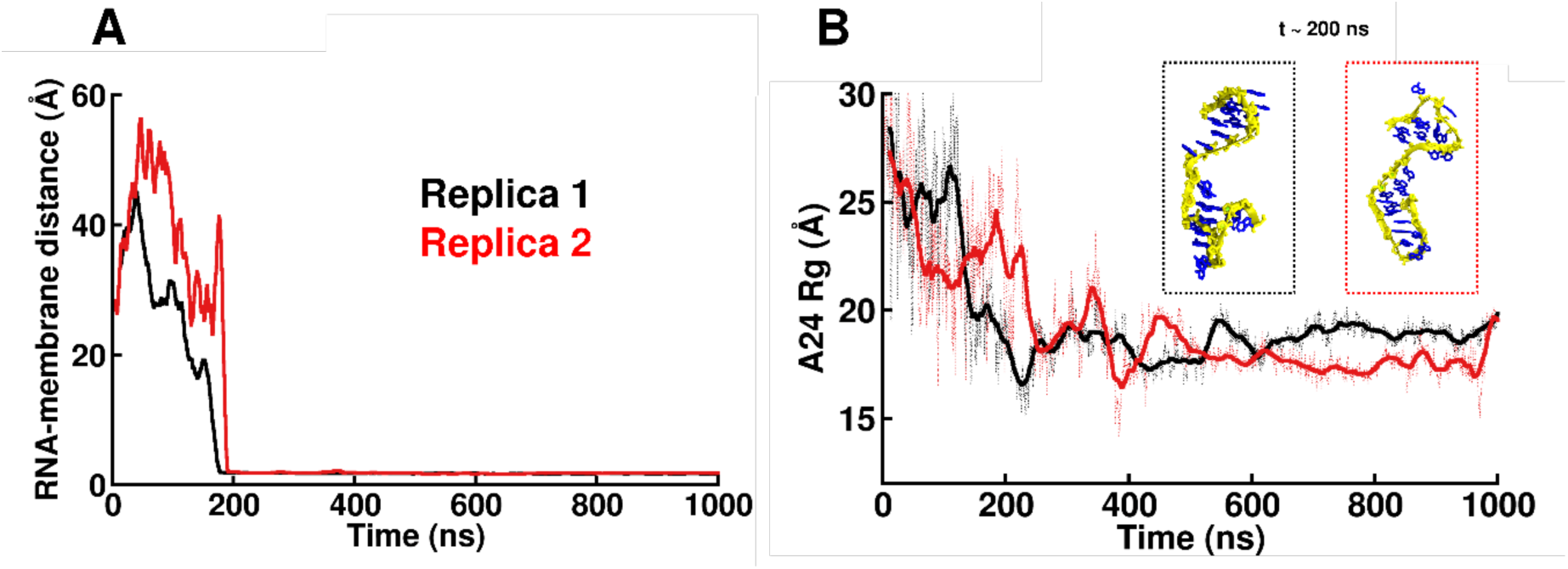
*(A)* Minimum distance between A24 and DPPC-gel bilayer. *(B)* Radius of gyration of A24 over the simulation time. Inset: structural arrangement of *A24* in both A24/DPPC-gel bilayer replicas at t = 200 ns. Fig S6A and Fig S6B show that *A24* took approximately 200 ns to fold through base-base stacking, which concomitantly led to adsorption to the DPPC-gel bilayer.

**Figure S7.**
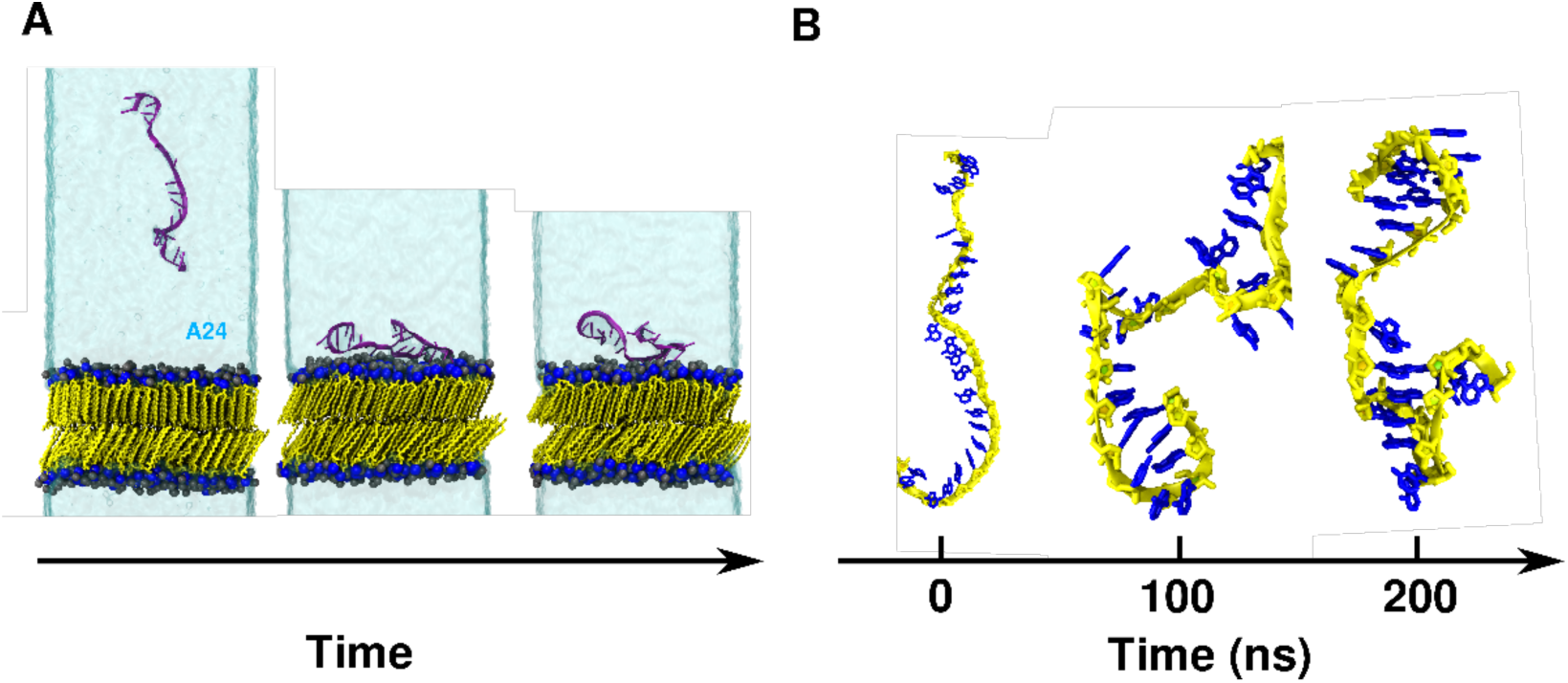
(A) Representative snapshots from MD simulations of the A24/DPPC-gel bilayer system. Color scheme: Blue and black for nitrogen and phosphate atom of lipid head groups, yellow for lipid tails, translucent ice blue for water. RNA structures are depicted in new-cartoon (magenta) representation. *(B)* Structural rearrangement of A24 in the A24/DPPC-gel bilayer system over time. Color scheme: yellow for A24 backbone and blue for nucleic bases.

**Figure S8.**
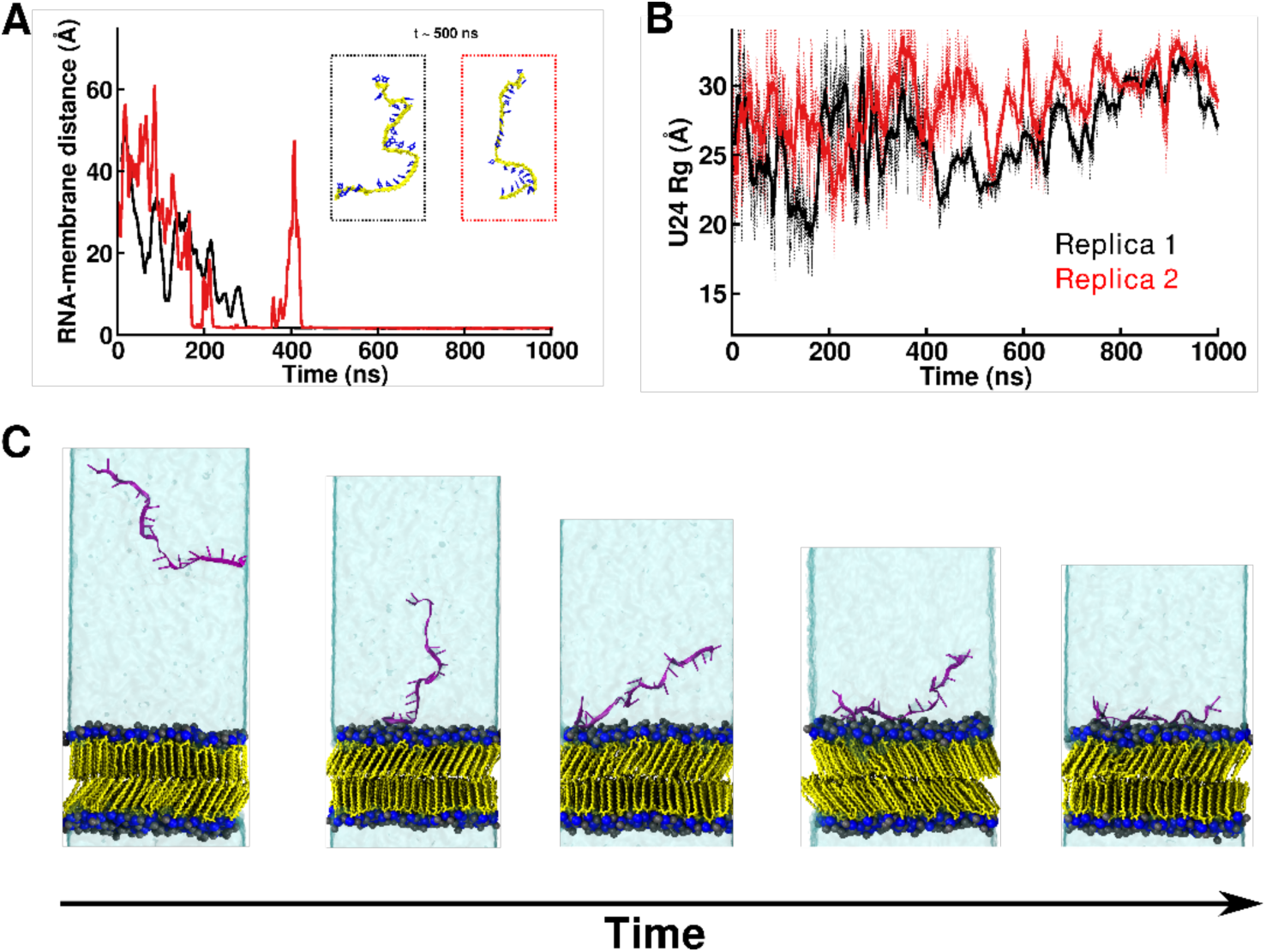
*(A)* Minimum distance between U24 and DPPC bilayer. Inset: Structural arrangement of *U24* in both U24/DPPC-gel bilayer replicas after adsorption (at t = 500 ns). *(B)* Radius of gyration of U24 over the simulation time. *(C)* Representative snapshots from MD simulations of the U24/DPPC-gel bilayer systems. Color scheme same as the Fig. S2.

**Figure S9.**
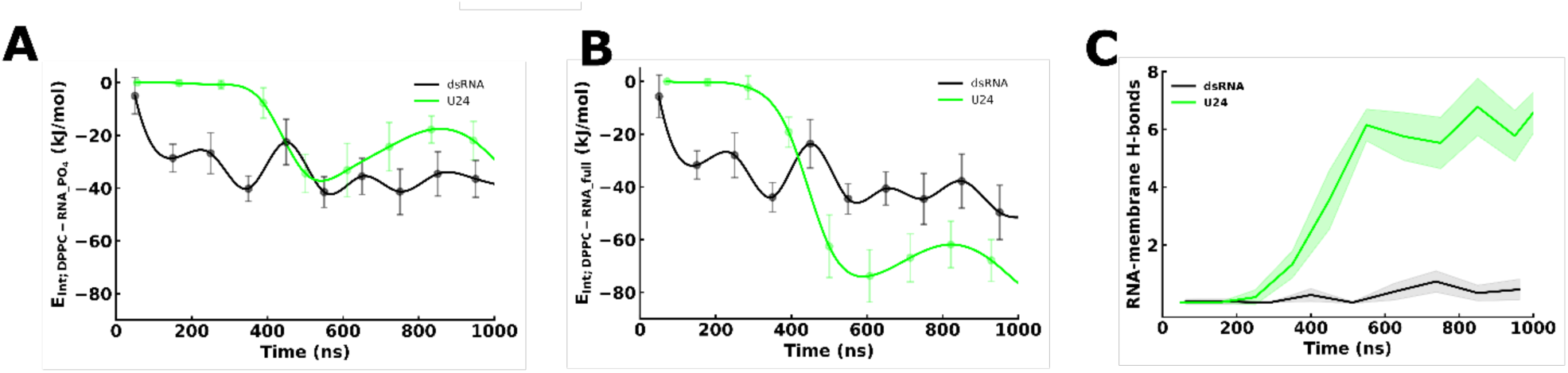
Interaction energy per nucleotide (*E_Coul._* + *E_LJ_*) over time. *(A)* phosphates 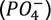 of dsRNA or U24 backbone with DPPC-gel bilayer, *(B)* All atoms of dsRNA or U24 with DPPC-gel bilayer. (C) DPPC-RNA H-bonds in dsRNA or ssRNA/DPPC-gel bilayer systems, as a function of time. Solid line indicates the running average, shade area indicates the standard deviation.

**Figure S10.**
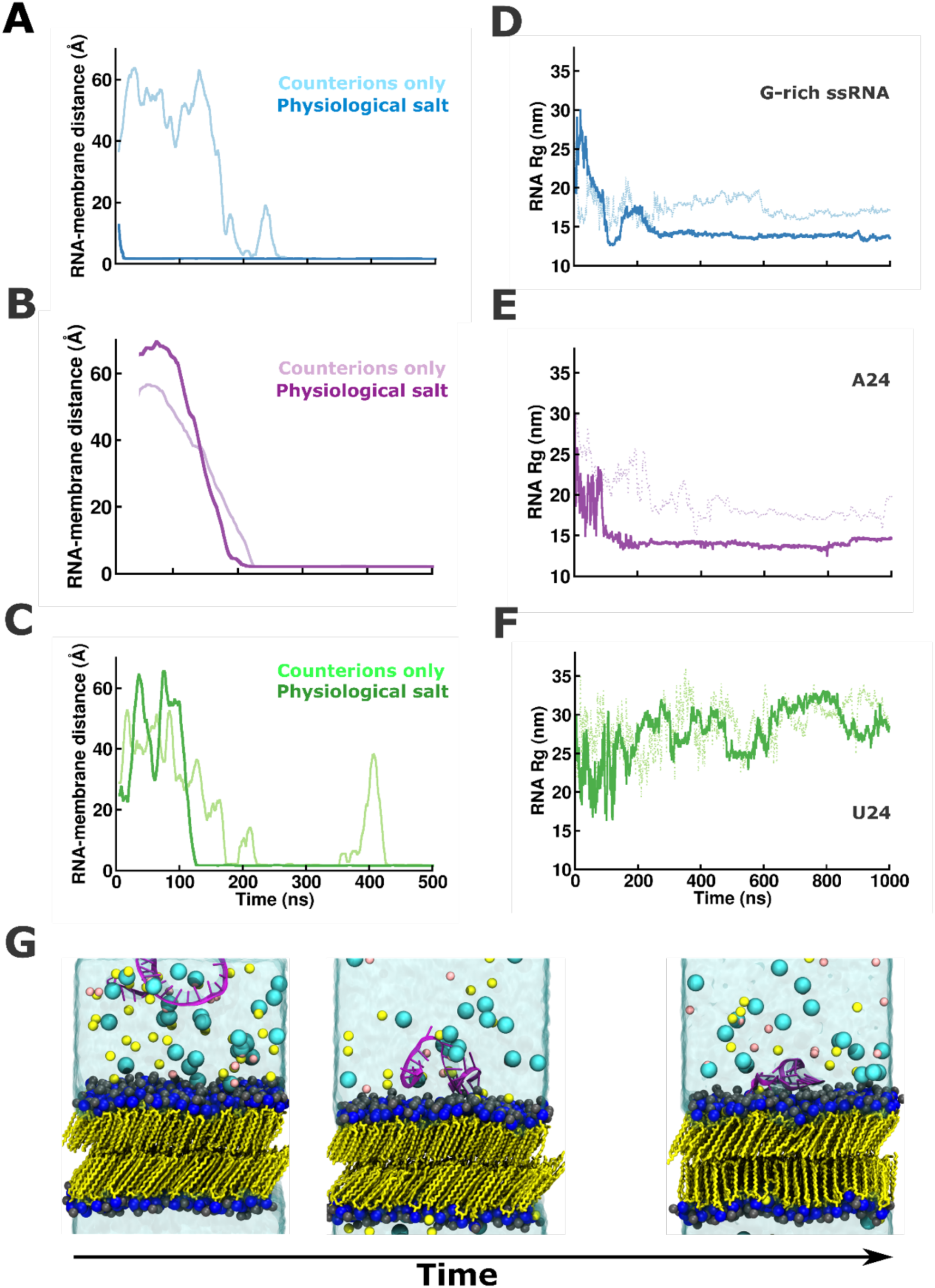
Comparison analysis of minimum distance between ssRNA and DPPC-gel bilayer *(A, B, C)*, and *corresponding* radius of gyration of ssRNA *(D, E, F)* in absence and presence of physiological salt (150mM NaCl + 50 mM MgCl_2_). *(C)* Representative snapshots from MD simulations of G-rich ssRNA/DPPC-gel bilayer systems in presence of physiological salt. Color scheme same as the Fig. S2A.

**Figure S11.**
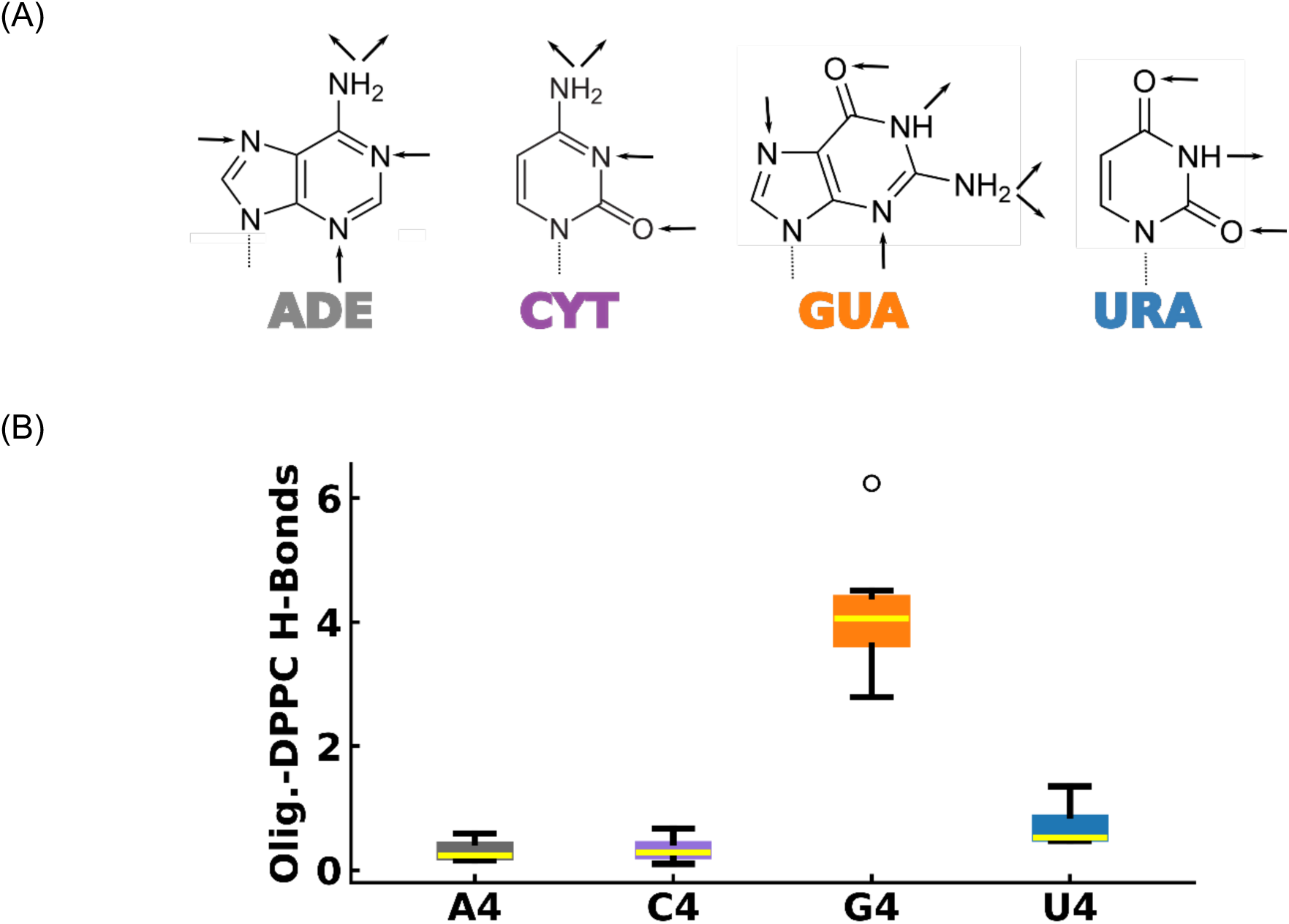
(A) Molecular structures of nucleic bases, showing the number of interaction sites (acceptor and donor atoms). Guanine has higher acceptor and donor H-atoms to form H-bonds. (B) MD simulation analysis of oligomers-DPPC bilayer H-bonds. Color scheme: gray for A4, purple for C4, orange for G4 and blue for U4 oligomers.

**Table S1.** Details about the systems simulated in this study.

| System name | Lipids per Leaflet/Total | Length of water slab<br>in Z-direction (Å) |
| --- | --- | --- |
| A4/DPPC-gel bilayer | 64/128 | 80 |
| C4/DPPC-gel bilayer | 64/128 | 80 |
| G4/DPPC-gel bilayer | 64/128 | 80 |
| U4/DPPC-gel bilayer | 64/128 | 80 |
| dsRNA/DPPC-gel bilayer | 144/288 | 180 |
| dsRNA/DOPC-liquid<br>bilayer | 144/288 | 180 |
| G-rich-ssRNA/DPPC-gel<br>bilayer | 144/288 | 180 |
| A24-ssRNA/DPPC-gel<br>bilayer | 144/288 | 180 |
| U24-ssRNA/DPPC-gel<br>bilayer | 144/288 | 180 |

All systems are neutralized by just $Na^+$ and $Cl^-$ ions or an explicit physiological salt concentration of 150 mM $NaCl$ + 50 mM $MgCl_2$ .

**Table S2.**
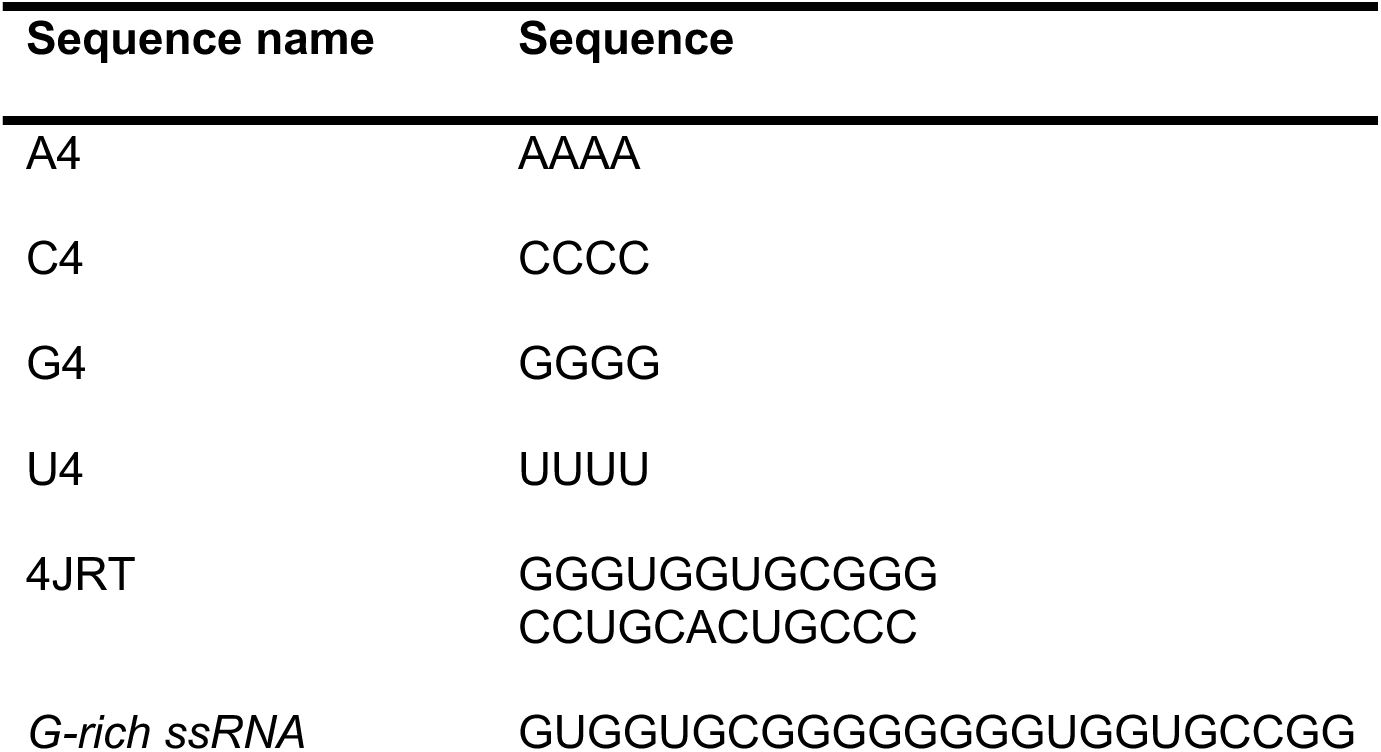
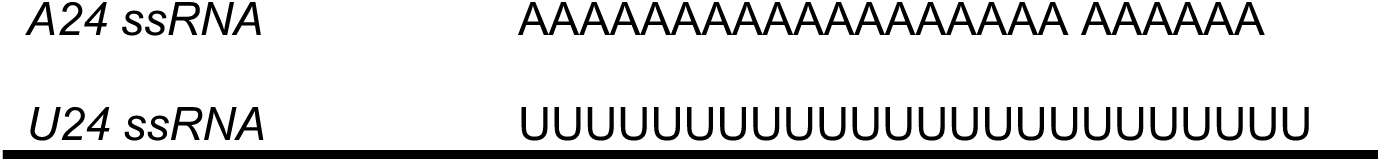
Details about RNA sequences used in the study.

### Supplementary Methods

#### Simulations Details

All MD simulations in this work were carried out with the simulation software GROMACS, version 2021.5.^1–3^ DPPC and DOPC lipid bilayer structures were built using the CHARMM-GUI package, ^4–5^ and then equilibrated for 50 ns at 298K and 310K, respectively, and 1 atm. The resulting configurations were used as the initial structure to build the final model systems: lipid bilayers in the presence of short-chain oligomers, either double-stranded RNA (dsRNA) or single-stranded RNA (ssRNA), through PACKMOL.^6^ Ions were placed randomly in the simulation box. Short-chain oligomers and ssRNA PDB structures were determined using the RNAComposer software.^7^ The dsRNA structure of 4JRT^8^ was taken from the Protein Data Bank (PDB). The *CHARMM36* forcefield was employed for all simulations.^9–12^ CHARMM TIP3P water model^13,14^ was used to solvate all systems. TopoTools, the virtual molecular dynamics (VMD) plugin, was used to convert PSFGEN generated topologies into GROMACS file formats.^15,16^ All simulations were performed with a 2 fs time step, and data was saved every 10 ps. The temperature (298 K or 310 K) was maintained with the Nosé−Hoover thermostat,^17,18^ and the semi-isotropic Parrinello−Rahman barostat was used to control the pressure.^19^ All the nonbonded interactions were truncated at a cutoff distance of 12 Å. The Particle Mesh Ewald (PME) method was used for the efficient computation of long-range electrostatic using a real space cutoff of 12 Å. Periodic boundary conditions (PBC) were applied in all directions.

All systems were minimized using steepest descent algorithm to eliminate any possible clashes and bad contacts. Subsequently, an NVT ensemble simulation was conducted for 2 ns using position restraints to all heavy atoms. Then, NPT ensemble simulations were performed for 5 ns for further equilibration. Finally, production runs were carried out for 1µs for all systems at the above described conditions, with no restraints. The simulation time have been chosen based on prior MD simulations studies ^20–24^, and to obtain convergence for multiple observables, including interaction energy (Figs S1, S9) and radius of gyration (Figs S5, S6, S8, S10). Selected snapshots were rendered using the VMD software package.

#### Binding free energy

The potential of mean force (PMF) for the G4 oligomer binding to the DPPC-gel bilayer as a function of distance was computed using the umbrella sampling algorithm^25^ and the weighted histogram analysis method (WHAM).^26^ The G4 oligomer was pulled from bound state to unbound state along the direction of the bilayer normal (z-axis) with a force constant of 1200 kJ mol^−1^ nm^−2^, for a total of 35 windows with a window spacing of 0.1 nm. The 35 windows were equilibrated for 1 ns and then used as starting configurations for umbrella sampling simulations. Each window was simulated for 500 ns. Error bars on the PMF curves are estimated using block averaging.

#### Interaction Energy

The interaction energy based on short-range energy components was calculated using GROMACS energy groups^27–31^ by decomposing the potential energy *via* rerunning the trajectories as follows:

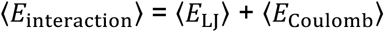

#### Hydrogen bond definition

We analyzed hydrogen bonds (H-bonds) between all possible donors and acceptors atoms with the GROMACS program *gmx hbond*. It characterizes an H-bond between a “donor” atom (a hydrogen connected to oxygen or other electronegative atom) and “acceptor” atom (oxygen or another electro-negative atoms) by two parameters: the radial distance between electronegative atoms, r, and the angle formed between the hydroxyl and the vector connecting the oxygen atoms, θ. To determine if an H-bond exists, a geometrical criterion was used: r is ≤ 3.5 Å and θ ≤ 30°.^32–34^

### Radius of gyration

We calculated the radius of gyration (*R_g_*) with the GROMACS program *gmx gyrate* as follows:

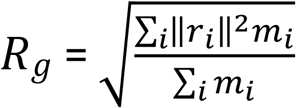

where *m_i_* and *r_i_* are the mass and the position of atom *i* with respect to the center of mass of the molecule, respectively.

### Cluster analysis and conformational entropy

Oligomer conformational entropy is associated with the number of its attainable conformational microstates and respective probabilities as such:

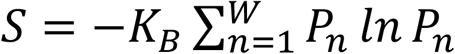

where *K_B_* is the Boltzmann constant, *W* is the number of unique conformational basins, and *P_n_* is the probability of being in conformational basin *n*. The unique conformations were determined using cluster analyses on MD trajectories performed by the GROMACS program *gmx cluster*. We used *gromos* algorithm for clustering.^35^ This method performs a clustering procedure based on the values of root-mean-squared deviation (RMSD). We choose the oligomer backbone for least squares fit and RMSD calculation with 0.15 nm cut-off value.

### Relative shape anisotropy

The relative shape anisotropy 𝜅^2^ is obtained from estimates of the radius of gyration *R_g_*, asphericity *b*, and acylindricity *c* as follows:

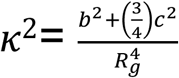

where *Rg* s the radius of gyration, *b* the asphericity, and ***c*** the acylindricity: 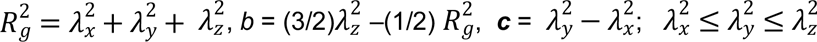, are the principle moments of the gyration tensor **S.**^36^

